# Integrated immunopeptidomics and transcriptomics reveal a cancer-restricted non-canonical HLA class I ligandome in medulloblastoma

**DOI:** 10.64898/2026.09.26.754718

**Authors:** Ningze Zheng, Yingqi Xu, Weihao mai, Aimin Hui

**Author notes:** Corresponding Author: Aimin Hui, Ph.D., State Key Laboratory of Respiratory Disease, Guangzhou Medical University, No. 195 Dongfeng West Road, Yuexiu District, Guangzhou 510182, Guangdong, China. These authors contributed equally (Ningze Zheng, Yingqi Xu, Weihao mai).

## Abstract

Medulloblastoma is the leading type of malignant brain tumor in children. Despite recent advances in T cell-based immunotherapy, the low mutational burden of medulloblastoma poses a considerable challenge to the discovery of mutation-derived neoantigens. Previous studies of medulloblastoma have investigated tumor-associated antigens and neoantigens derived from aberrant splice junctions. The landscape of non-canonical HLA class I-bound peptides (ncHLAp) remains poorly defined. To broaden the repertoire of targetable antigens in medulloblastoma, we mapped the ncHLAp landscape across the four major molecular subgroups of medulloblastoma (Group 3, Group 4, WNT, and SHH) by integrating immunopeptidomic and transcriptomic analyses. Candidates were further filtered against normal-tissue immunopeptidomic and ribosome profiling datasets to identify cancer-restricted ncHLAp (CR ncHLAp). A total of 2,298 ncHLAp were identified across 21 human medulloblastoma samples, of which 598 were CR ncHLAp. These CR ncHLAp were primarily derived from non-canonical translational sources, including retained introns, out-of-frame open reading frames, and long non-coding RNAs. Some CR ncHLAp were shared across medulloblastoma samples and detected in other cancer types. More than 70% of the CR ncHLAp were predicted to bind at least one patient-matched HLA allotype, and approximately 87% were predicted to have immunogenic potential. Collectively, this study defines a previously undercharacterized component, CR ncHLAp, of the medulloblastoma HLA-I ligandome and suggests that CR ncHLAp may serve as promising candidate targets for T-cell-based immunotherapy in medulloblastoma.

## Introduction

Medulloblastoma represents the most prevalent malignant brain tumor in children, accounting for approximately 20% of all pediatric central nervous system tumors [34]. According to distinct molecular features, medulloblastoma was classified into four subtypes, including group 3 (G3), group 4 (G4), WNT group, and SHH group [41]. Significant differences between the four subgroups are presented in DNA copy-number, transcriptome, and patient survival [21]. The current standard treatment for medulloblastoma consists of surgical resection followed by adjuvant radiotherapy and chemotherapy. However, radiotherapy can cause central nervous system damage, while limited drug penetration into the tumor site and the development of drug resistance remain major obstacles to chemotherapy. In addition, both radiotherapy and chemotherapy may lead to long-term, life-altering cognitive deficits in survivors [45]. Therefore, the development of new therapeutics for medulloblastoma with limited toxicity remains an unmet clinical need.

T cell-based immunotherapies offer a promising means of targeting malignant cells through recognizing epitopes presented by human leukocyte antigen class I (HLA-I) molecules. Immunotherapeutic strategies, including immune checkpoint inhibitors, oncolytic viral therapy, vaccine-based therapy, and adoptive cell therapy, have been investigated for the treatment of medulloblastoma [45]. Although these approaches have not yet produced consistent or robust clinical benefits in patients with medulloblastoma, preclinical studies have suggested their potential to enhance antitumor immune responses, supporting their continued investigation.

Therapeutic cancer vaccines are designed to elicit tumor-specific T-cell responses, enabling T cells to recognize and eliminate tumor cells. In recent years, the success of COVID-19 mRNA vaccines has accelerated the development of therapeutic mRNA cancer vaccines. An increasing number of these vaccines, targeting tumor-associated antigens (TAAs) or neoantigen-based tumor-specific antigens (TSAs), have entered clinical trials [48]. Although TAAs and mutation-derived neoantigens are abundant in many cancers and can be readily predicted, most TAAs and SNV- or INDEL-derived neoantigens are poorly immunogenic and provide limited clinical benefit to cancer patients [50]. This is particularly relevant to medulloblastoma, which has a low mutational burden [42]. Therefore, mutation-derived neoantigens may not represent ideal therapeutic targets for immunotherapy.

Nevertheless, emerging evidence showed that some peptides, called non-canonical antigens, are translated from noncoding regions, such as introns, long noncoding RNAs (lncRNAs), and pseudogene, and are presented on HLA-I [23, 29, 33]. Tumor-specific non-canonical antigens are absent from normal tissues and exhibit greater dissimilarity from self-antigens, making them more likely to elicit immunogenic responses [50]. Although the clinical efficacy of targeting non-canonical antigens remained to be established, preclinical studies have demonstrated that vaccination with these antigens induced robust CD8^+^ T-cells responses [1, 13, 24]. Unlike mutation-derived neoantigens, which are typically patient-specific, some non-canonical immunopeptides were found to be shared within same tumor and even across multiple cancers, thereby enabling the development of off-the-shelf cancer vaccines [46]. Moreover, cancer-restricted non-canonical antigens presented by HLA molecules on the cell surface can be directly identified using liquid chromatography-tandem mass spectrometry (LC-MS/MS), thereby expanding the repertoire of targetable antigens for cancer vaccine development [1, 13]. Although recently published studies have identified the TAAs and neoantigens originated from aberrant splice junctions, SNVs, and gene fusions in medulloblastoma [37, 42], the non-canonical antigens in this malignancy warrant further investigation.

In this study, we explored non-canonical antigens in medulloblastoma by reanalyzing published RNA-seq and HLA-I immunoprecipitation mass spectrometry datasets through integrated transcriptomic and immunopeptidomic analyses. We postulated that the identification of non-canonical antigens might reveal additional targetable antigens in medulloblastoma. To obtain a broader and more reliable repertoire of non-canonical antigens, we employed a two-step filtering approach during database searches of mass spectrometry data. Cancer-restricted non-canonical peptides (CR ncHLAp) were defined by excluding candidates with evidence of translation or HLA-I presentation in healthy tissues, as determined by Ribo-seq or HLA-I immunopeptidomics datasets. In addition, we predicted the foreignness and immunogenicity of CR ncHLAp using *in silico* algorithms and assessed whether these antigens were shared within individual cancer types and across cancer types. Finally, we used rMATS to identify aberrant alternative splicing (AS) events and investigate their potential association with the generation of CR ncHLAp in medulloblastoma.

## Methods

### Acquisition of raw sequencing and mass spectrometry data

The RNA-seq data of 21 medulloblastoma samples, including 6 samples from G3 subgroup, 6 samples from G4 subgroup, 3 samples from WNT subgroup, and 6 samples from SHH subgroup, were available from Gene Expression Ominibus (GEO) database with entries GSE238195 [42]. The mass spectrometry data of corresponding analyzed medulloblastoma samples were obtained from Proteomics Identifications Database (PRIDE) with entries PXD043770 [42]. For translation-centric filtering pipeline, we obtained the human normal tissues ribosome profiling data of 49 samples from 8 tissues from GEO database with accession codes GSE182377 and GSE59820 [9, 27]. In addition, HLA-I immunopeptidomics data from healthy tissues (59 samples across 29 tissues), including normal thymus samples but no testis samples, were downloaded from PRIDE under the dataset identifier PXD019643[30].

### Preprocessing of raw sequencing data

Files of raw RNA-seq data were converted to FASTQ format using the SRA Toolkit (RRID:SCR_024350) and were subsequently quality assessed by FastQC software (RRID:SCR_014583). FASTQ files that passed the quality control were processed with fastp (RRID:SCR_016962), including trimming sequencing adapters and quality filtering, ensuring a minimum read length of 30 bp after filtering. For each sample with multiple sequencing runs, trimmed R1 and trimmed R2 FASTQ files were separately merged to construct a single sample-level R1 and R2 FASTQ files, which further quality assessed using FastQC. The merged FASTQ files were then subjected to transcript expression quantification with kallisto (RRID:SCR_016582). In parallel, paired-end RNA-seq reads were aligned to the GRCh38 reference by using STAR (RRID:SCR_004463) with the following parameters: --alignSJoverhangMin 10, -- alignMatesGapMax 200000, --alignIntronMax 200000, and -- alignSJstitchMismatchNmax 5 -1 5 5. Indices for the BAM files were generated using SAMtools (RRID:SCR_002105). Single nucleotide variants (SNVs) with alternate allele count no less than 5 were called using freeBayes (RRID:SCR_010761). Biallelic SNVs with QUAL > 20 were retained after filtering with bcftools (RRID:SCR_005227) and were exported in a VCF file.

### Personalized protein databases generation for MS-based proteomic analysis

#### Personalized canonical proteome

To construct sample-specific personalized canonical proteome databases, biallelic SNVs identified from the corresponding RNA-seq data were incorporated into the GRCh38 reference genome using bcftools consensus. Based on the GRCh38 GTF annotation, canonical CDS sequences were extracted from the personalized genomes and translated into protein sequences using gffread (RRID:SCR_018965). Protein sequences retained after annotation-based filtering were concatenated to generate sample-specific personalized canonical proteome FASTA files.

#### Personalized retained introns proteome

For construction of personalized retained intron-derived protein databases, reference transcript sequences were first generated from the GRCh38 reference genome and Ensembl gene annotation using gffread. Intron pseudo-transcripts were then generated based on the GRCh38 reference genome and Ensembl gene annotation. In the intron pseudo-transcripts, intronic sequences were extended 25 bp into the adjacent exons at both ends. An augmented transcriptome was constructed by combining the reference transcriptome with the intron pseudo-transcripts. Using kallisto, merged RNA-seq FASTQ files were pseudoaligned to the augmented transcriptome, and the expression levels of annotated transcripts and intron pseudo-transcripts were quantified. Retained introns were called using defined parameters: intron TPM ≥ 1, exon TPM ≥ 1, intron retention ratio ≥ 0.05, and at least 5 unique intron-supporting reads. Subsequently, biallelic SNVs from the VCF files generated in the previous section were incorporated into the retained-intron coding windows. Finally, both reference and variant-recoded retained-intron sequences were translated in three reading frames. Amino acid sequences with a length of at least 8 residues were retained and concatenated to construct retained intron-derived protein FASTA databases.

Moreover, IRFinder was employed to identify additional retained introns. In this pipeline, coordinate-sorted BAM files generated in the previous section were first converted to name-sorted BAM files using SAMtools. IRFinder was then run in BAM mode. Retained introns were kept if they met the defined criteria: intron coverage > 0.5, intron depth > 3, exon-to-intron reads at both left and right junctions > 2, exact splice reads > 4, and intron retention ratio > 0.03. The filtered retained introns were converted into coding intron sequences using the GRCh38 genome and Ensembl GRCh38.88 GTF annotation as references. The coding intron sequences were translated in three reading frames, and the resulting amino acid sequences with a length of at least 8 residues were retained. Finally, the resulting sequence were reassembled into IRFinder-derived intron protein FASTA database.

#### Personalized novel or unannotated open reading frame (nuORF) proteome

The nucleotide sequences of reference nuORFs were extracted from the GRCh38 toplevel genome according to the nuORFdb_v1.2 BED file obtained from the Broad Institute website [33]. Subsequently, sample-specific RNA-seq-derived biallelic SNVs identified in the previous section were incorporated into the corresponding reference nuORF sequences using the ALT allele. Nucleotide sequences of both reference nuORFs and variant-containing nuORFs were translated in three reading frames. Finally, the resulting amino acid sequences with a length of at least 8 residues were compiled into nuORF-derived protein databases.

#### Personalized small open reading frame (smORF) proteome

A curated smORF database comprising lncRNA-derived smORFs and uORFs was obtained from a previous study by Prensner et al [35]. To construct patient-specific smORF-derived protein databases, smORF annotations were extracted from this database. Reference nucleotide sequences were retrieved from the GRCh38 primary assembly genome according to the genomic coordinates provided in the annotations. For each sample, biallelic SNVs identified by FreeBayes were incorporated into the corresponding smORF sequences when available. The resulting sample-specific smORF nucleotide sequences were then translated into amino acid sequences. Protein sequences with a minimum length of 8 amino acids were retained and aggregated to generate personalized smORF-derived protein databases.

#### Personalized gene fusion proteome

Gene fusions were identified from RNA-seq data using Arriba v2.5.1 (RRID:SCR_025854). The reference genome was indexed using STAR with the GRCh38 genome FASTA and GENCODE v49 GTF file. The merged and trimmed paired-end RNA-seq FASTQ files for each sample, generated in the previous section, were aligned to the reference genome using STAR with chimeric alignment enabled. The resulting coordinate-sorted BAM files were subjected to Arriba for gene fusion detection. Predicted gene fusion events were further filtered by retaining events with high or medium confidence, in-frame or out-of-frame reading frames, and a total supporting read count of at least 2. Junction-centered fusion sequences were generated by extracting 30 amino acids upstream and 30 amino acids downstream of the fusion breakpoint. The resulting junction 60-aa sequences were aggregated into personalized gene fusion-derived protein databases.

#### Combined personalized protein databases

To define the search space for LC-MS/MS data analysis, FASTA headers in the personalized protein databases and the common laboratory contaminant database were appended with source-specific PE tags following the FragPipe-compatible format. The contaminant database was downloaded from The Global Proteome Machine (GPM) website. For each patient sample, the personalized canonical proteome, together with contaminant protein sequences, was integrated separately with retained intron-derived, IRFinder-derived intronic, nuORF-derived, smORF-derived, or gene fusion-derived protein sequences to generate source-specific search databases. In parallel, a fully combined search database was constructed by merging the personalized canonical proteome and contaminant proteins with all non-canonical protein databases.

### MS/MS search processes

All raw mass spectrometry data were first converted to mzML format using ProteoWizard (RRID:SCR_012056) with the peak-picking mode enabled to ensure compatibility with FragPipe (RRID:SCR_022864). To improve the accurate identification of non-canonical peptides, a two-step database-searching strategy was applied. In the first step, LC-MS/MS data from each patient sample were searched separately against the corresponding personalized protein databases, including personalized canonical, retained intron-derived, IRFinder-derived intronic, nuORF-derived, smORF-derived, gene fusion-derived, and fully combined protein databases. Database searches were performed using the nonspecific HLA peptidomics mode with the following parameters: peptide length of 7-15 amino acids, decoy sequences appended to the search database, and a group-level PSM false discovery rate (FDR) threshold of 5%. For each sample, all peptides identified in the first-step search were aggregated to construct a reduced search database. In the second step, the original LC-MS/MS data were re-searched against the corresponding reduced database with a group-level PSM FDR threshold of 1%, while the other parameters were kept the same as those used in the first step.

### Normal tissues Ribo-seq data process and used for HLA-I-bound peptides (HLAp) filtering

To remove MS-identified peptides with translational evidence in healthy tissues, normal-tissue Ribo-seq data were preprocessed and used for peptide filtering. A customized transcriptome annotation was generated by integrating GENCODE v26 annotation, CHESS 3.1.1 GRCh38 primary annotation, and lincRNA transcripts from Cabili et al [3]. The resulting custom GTF annotation was used for STAR index generation and RiboTISH-based translated ORF prediction. Each FASTQ file was trimmed using Trimmomatic v0.36 (RRID:SCR_011848) in single-end mode. The trimmed reads were aligned to contaminant RNA sequences using Bowtie2 (RRID:SCR_016368). The reads unmapped to contaminant RNA sequences were retained and further aligned to the GRCh38 reference genome using STAR with the following parameters: --alignSJDBoverhangMin 1, --alignSJoverhangMin 51, -- outFilterMismatchNmax 2, --alignEndsType EndToEnd, --alignIntronMin 20, -- alignIntronMax 100000, --outFilterType BySJout, --outFilterMismatchNoverLmax 0.04, --twopassMode Basic, and --outSAMattributes MD NH. The resulting SAM files were converted to BAM format. For each tissue, BAM files from corresponding samples were merged and subsequently subjected to quality assessment and translated ORF prediction using RiboTISH. Predicted ORFs were retained if they met both criteria: RiboPvalue < 0.05 and FrameQvalue < 0.05. MS-identified peptides from each sample were then matched against normal-tissue translated ORF amino acid sequences. During sequence matching, isoleucine and leucine were not distinguished. Finally, peptides found in any healthy-tissue translated ORF were removed.

### Normal tissues immunopeptidome data reprocess and used for non-canonical HLAp (ncHLAp) filtering

Raw mass spectrometry data from normal tissues, including 54 samples from 28 healthy tissues and 5 samples from normal thymus, were downloaded and reanalyzed using FragPipe. For healthy tissues, the immunopeptidomics data were searched separately against the personalized proteome databases generated for the 21 medulloblastoma patients, using the same search workflow and parameters as those applied to the medulloblastoma samples. Peptides with a probability score ≥ 0.9 and a spectral count ≥ 2 were extracted to generate an exclusion list. Ribo-seq-filtered candidate peptides were removed if their sequences exactly matched any peptide in the exclusion list, with isoleucine and leucine treated as equivalent. Similarly, the normal thymus MS data were reprocessed using the same workflow and parameters. Peptides were included in the thymus-derived exclusion set if they met the following criteria: probability score ≥ 0.9; a spectral count ≥ 2 in at least one thymus sample or spectral counts > 0 in at least two thymus samples. Candidate peptides retained after healthy tissue filtering were further discarded if they exactly matched any peptide in the thymus-derived exclusion set.

### HLA allele prediction of HLAp

HLA allele-binding predictions were performed for MS-identified peptides across the 21 medulloblastoma samples using HLAthena (RRID:SCR_028691). The HLA alleles for each patient were obtained from the study by Julia et al [42]. To compare the amino acid sequence profiles of canonical HLAp and ncHLAp predicted to bind a given HLA allele, 9-mer peptides with a predicted percentile rank ≤ 2% for the corresponding HLA allele were selected and used to generate peptide sequence profile plots using MHCMotifDecon [19].

### Code Availability Statement

The code used in this study was adapted from Zack-Ely/PDAC-ncHLAp-Project (version 1.0; https://zenodo.org/records/14743979) [13] and is available in Code Ocean at https://doi.org/10.24433/CO.4112889.v1.

## Results

### Detection of ncHLAp in medulloblastoma immunopeptidome

Previous studies querying the medulloblastoma immunopeptidome have primarily focused on TAAs and TSAs derived from SNVs, gene fusions, and aberrant splice junctions [37, 42]. However, development of T cell-based immunotherapy for medulloblastoma heightens the need to look beyond to investigate non-canonical peptides presented on HLA-I molecule. Therefore, we investigated non-canonical peptides of medulloblastoma by integrating transcriptome and immunopeptidome.

In the present work, we obtained HLA-I immunoprecipitation-MS data and paired RNAseq data of 21 patients from the study of Julia Velz et al [42]. The mean purity of the 21 tumors is 84.2% (Supplementary Fig. 1A). RNA sequencing data were used for variant calling and kallisto quantification, which were then subjected to personalized protein databases construction for each patient. To improve the accuracy of non-canonical peptide identification, MS raw data were separately searched against these custom MS databases using FragPipe with two-step filtering strategy. In the first step, the search was performed at group-level 5% FDR, and the filtered peptides were assembled into a reduced protein database for each sample. In the second step, the MS raw data were re-searched against the reduced protein database with group-level 1% FDR. Finally, the retained candidates were annotated as canonical or non-canonical peptides (Fig. 1A).

**Fig. 1.**
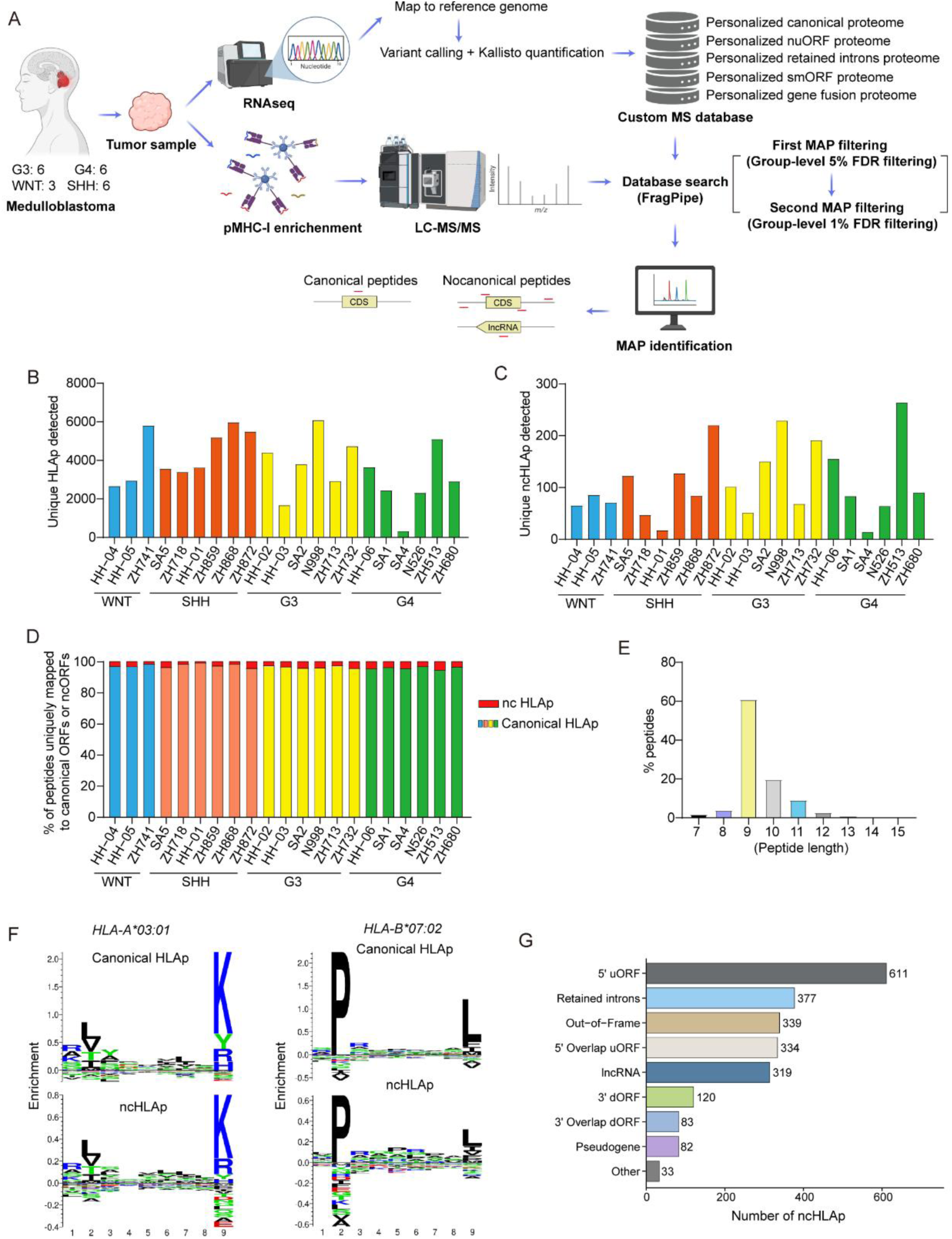
Immunopeptidomics approach to identify ncHLAp in medulloblastoma. **(A)** Workflow depicting the non-canonical antigens discovery strategy. The RNA sequencing data and mass spectrometry data of medulloblastoma samples were generated by Julia Velz *et al*. Proteomic spectra were searched against custom databases, which were created based on RNA-seq, under two-steps FDR filtering. The MHC I-associated peptides (MAP) were identified and annotated as canonical or nocanonical peptides. Schematic diagrams were created in BioRender. Number of HLAp **(B)** or ncHLAp **(C)** uniquely detected in each medulloblastoma samples, which were grouped based on molecular subgroup (WNT, SHH, G3, and G4). **(D)** Proportion of canonical ORFs derived HLAp or non-canonical ORFs (ncORFs) derived HLAp per medulloblastoma sample. **(E)** Length distribution of HLAp across all the samples. **(F)** Comparison of the amino acid profile of 9-mer canonical HLAp and ncHLAp predicted to bind HLA-A*03:01 or HLA-B*07:02. HLA allele binding was predicted by HLAthena and peptide sequence profiles were plotted using MHCMotifDecon. **(G)** Number of ncHLAp originated from different nuORF biotypes in all analyzed samples.

Using the above two-step FDR filtering strategy, we identified a median of 3622 [330–6080] distinct HLAp and 85 [14–264] unique ncHLAp per sample (Fig. 1B and Fig. 1C). ncHLAp accounted for 0.47% to 5.2% (median, 3.1%) of the overall HLAp in each patient tumor (Fig. 1D). In line with previous studies, most HLAp were typically 8- to 11-mers (Fig. 1E) [13, 42]. An enrichment for canonical HLAp and ncHLAp predicted to bind to HLA-A*02*01, HLA-A*03*01, HLA-A*11*01, HLA-A*68*01, and HLA-B*07*02 was observed (Supplementary Fig. 1B). Comparative profiling reflected that sequence motifs of canonical HLAp and ncHLAp predicted to bind to HLA-A*03*01 and HLA-B*07*02, two representative allotypes, were no obvious different (Fig. 1F). By annotating the gene biotypes of ncHLAps across the 21 samples, we found that peptides derived from 5′ upstream ORFs (uORFs) were overrepresented, which is consistent with the findings in other tumor types [13]. Besides, some ncHLAp were found to be produced by translation of retained introns, Out-of-frame, 5’ overlap uORF, and lncRNA (Fig. 1G). Together, these observations indicated that a substantial number of ncHLAp were identified in medulloblastoma by using immunopeptidomics.

### Identification and characterization of cancer-restricted non-canonical HLA class I-bound peptides (CR ncHLAp) in medulloblastoma

Previous studies have demonstrated that substantial non-canonical peptides show evidence of translation and are presented by HLA-I in healthy tissues [13, 30]. Moreover, the T cells strongly recognized self peptide-MHC complexes undergo negative selection in thymic medulla [20]. Therefore, it is necessary to identify cancer restricted ncHLAp, nominating therapeutic targets. In this study, we assessed the tumor specificity of ncHLAp according to the previously described method [13]. The Ribo-seq data of 49 samples from 8 distinct healthy tissues [9], immunopeptidomics of 54 samples from 28 different normal tissues [30], and immunopeptidomic data of 5 samples from healthy thymus tissues were leveraged in the filtering process (Fig. 2A) [30]. Following selection, 598 of the 2298 ncHLAp identified above were found to be tumor-specific, with a median of 26 [5–72] CR ncHLAp per patient specimen (Fig. 2B). The identities of representative CR ncHLAp were supported by their MS/MS fragmentation spectra (Supplementary Fig. 2A-C). Meanwhile, canonical HLAp detected in the same 21 medulloblastoma samples were analyzed in parallel using the same filtering procedure applied to candidate CR ncHLAp. As expected, the source ORFs of 99.04% of the canonical HLAp showed evidence of translation in at least one healthy tissue (Fig. 2C), in line with previous reports [13, 22, 30]. This result supports the ability of our filtering pipeline to identify and exclude HLAp whose source ORFs are translated in healthy tissues or that are detected in healthy-tissue immunopeptidomes. Consistent with the known HLA-I peptide-binding preferences, the majority of CR ncHLAp were 8-11 amino acids long (Fig. 2D). Compared with non-CR ncHLAp, CR ncHLAp exhibited a distinctive biotype distribution, with a large proportion of CR ncHLAp derived from translation of retained introns, suggesting that introns were preferentially retained and translated in medulloblastoma (Fig. 2E). To determine whether CR ncHLAp displayed any altered physical properties, we compared the calculated hydrophobicity index (HI) of CR ncHLAp and canonical HLAp against the observed retention time (RT). The RT distribution of CR ncHLAp showed no significant difference from that of canonical HLAp, indicating their correct identification (Fig. 2F). However, the relative foreignness of CR ncHLAp was predicted to be more obvious than that of canonical HLAp (Fig. 2G). HLA-I affinity analysis for CR ncHLAp-MHC pairs indicated that 73.3% of CR ncHLAp were predicted to bind to their cognate HLA, with over 50% showed high MHC I-binding affinity (Fig. 2H). Immunogenicity prediction also indicated that over 87% CR ncHLAp with 9 or 10 amino acids in length exhibited marked immunogenicity (Fig. 2I). Overall, these data suggest that the CR ncHLAp identified in medulloblastoma served as potential therapeutic targets.

**Fig. 2.**
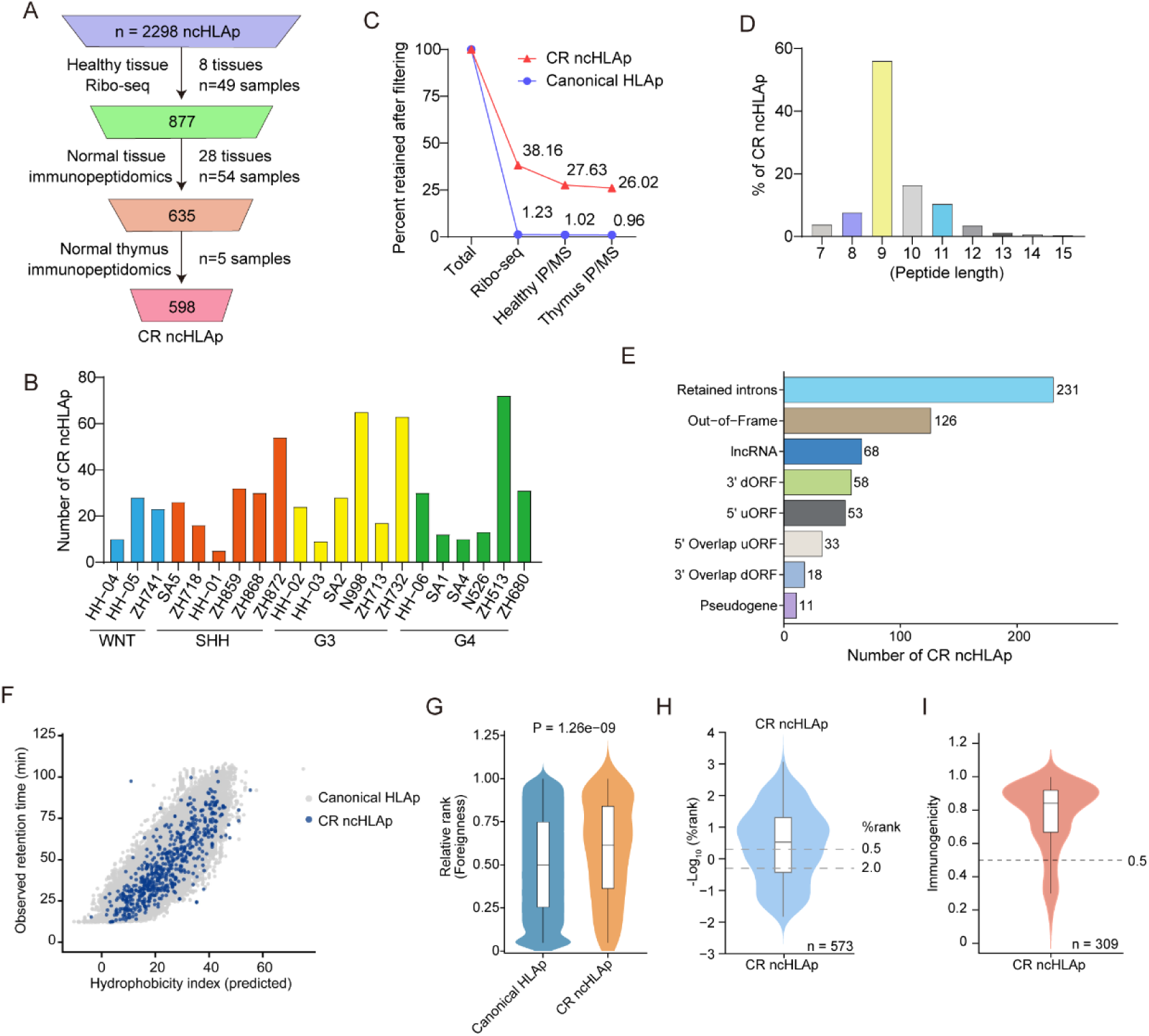
Filtering and characterization of CR ncHLAp in medulloblastoma. **(A)** A translation-centric filtering pipeline was employed to exclude ncHLAp whose source ORFs showed evidence of translation in any of the normal tissues examined or that were detected in normal-tissue HLA-I immunopeptidomes. CR ncHLAp across all samples were retained following the filtering steps. **(B)** The number of CR ncHLAp in each medulloblastoma sample. **(C)** Percentages of retained CR ncHLAp and canonical HLAp following each step of the filtering pipeline. **(D)** Length distribution of CR ncHLAp in all the samples. **(E)** The number of CR ncHLAp derived from different nuORF biotypes across all patient samples. **(F)** The hydrophobicity index was plotted against observed retention time for identified canonical HLAp and CR ncHLAp. Hydrophobicity index was predicted locally using an SSRCalc-like sequence-specific retention calculator implemented in R, based on the Krokhin model. **(G)** Comparison of the relative foreignness of canonical HLAp and CR ncHLAp. **(H)** Boxplots of the HLA-I affinity for all CR ncHLAp-MHC pairs. The affinity was predicted by NetMHCPan 4.1 and was presented as rank order. **(I)** Immunogenicity prediction by deepimmno for the identified 9-mer and 10-mer CR ncHLAp [25].

### A subset of CR ncHLAp are shared among medulloblastoma samples and across different tumor types

While mutation-derived neoepitopes were predominantly patient-specific, many CR ncHLAp were found to be shared among medulloblastoma samples. Given that HLA-A*11:01 was one of the most common HLA allotypes in our cohort, we assessed the recurrence of CR ncHLAp predicted to bind this allotype. Approximately 40% of these peptides were shared by two or more HLA-A11:01-positive patients with medulloblastoma (Fig. 3A). Although the 598 CR ncHLAp described above were classified as cancer-restricted using our translation-centric filtering approach, it remained unclear whether their source transcripts also exhibited tumor-restricted expression. To address this question, we retrieved normal-tissue expression data from the Developmental Genotype-Tissue Expression (dGTEx) project for the source transcripts of the 30 CR ncHLAp with the highest transcript abundance in medulloblastoma, including recurrent CR ncHLAp shared across patient samples. Consistent with a previous study, the source transcripts of a subset of CR ncHLAp were detected at only trace levels or remained undetected in healthy tissues; however, most source transcripts that were highly expressed in medulloblastoma were also expressed at appreciable levels in normal tissues (Fig. 3B). This finding suggests that the generation of CR ncHLAp may result from tumor-associated alterations in RNA processing or translation of their source transcripts. We next investigated the recurrence of the CR ncHLAp identified in this study across the 15 cancer types included in the IEAtlas dataset [4]. Our results showed that more than 30 CR ncHLAp were detected in at least one additional cancer type, providing independent evidence for their HLA-I presentation and reducing the likelihood that these identifications resulted from analytical artifacts (Fig. 3C). Finally, selected CR ncHLAp were validated by PRM and a representative MS/MS spectrum is shown in Fig. 3D. Taken together, these findings suggest that a subset of CR ncHLAp recur across medulloblastoma samples and other cancer types and may represent promising targets for T cell-based cancer immunotherapy.

**Fig. 3.**
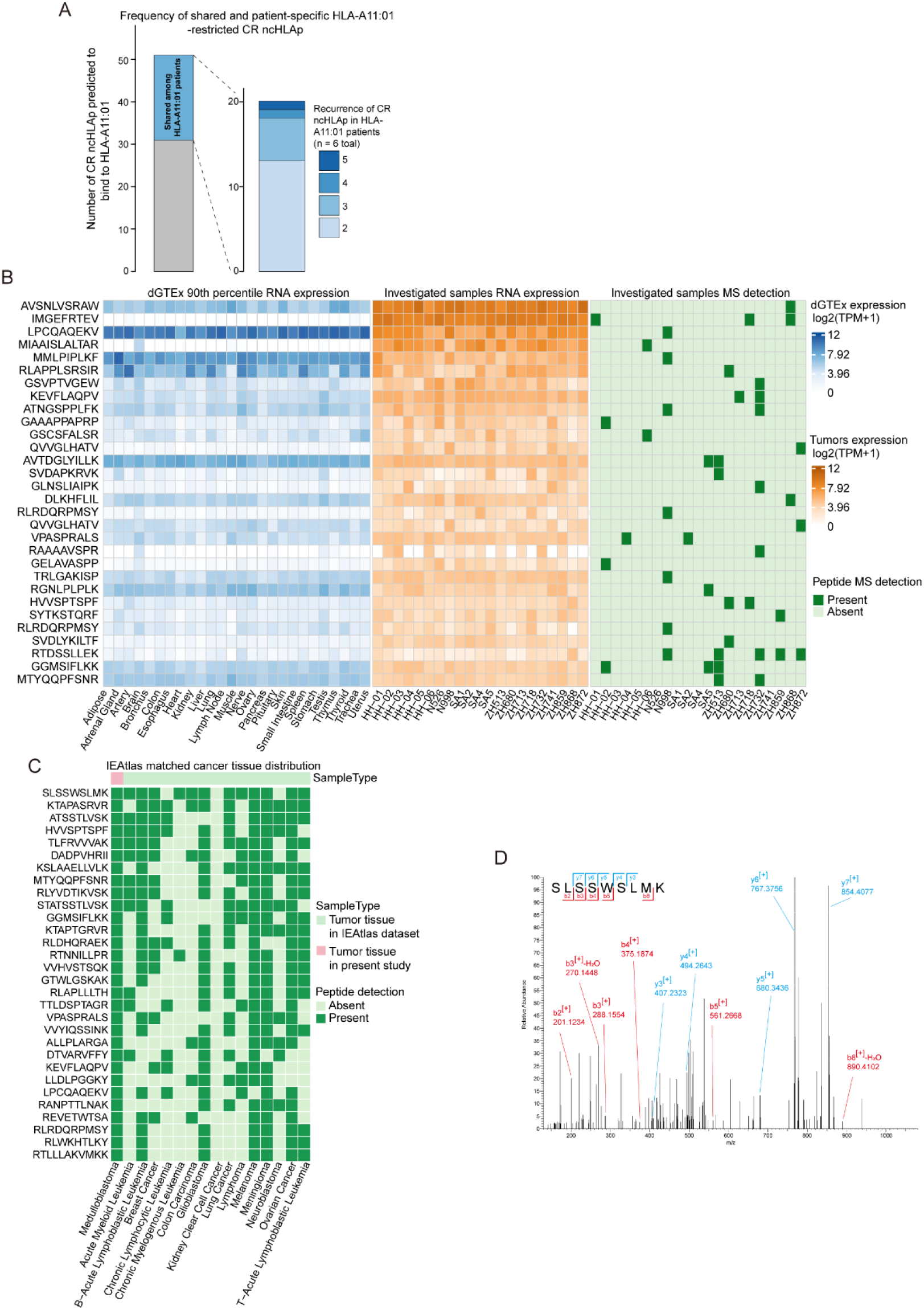
CR ncHLAp are shared across patients with medulloblastoma and across different tumor types. **(A)** Numbers of CR ncHLAp predicted to bind to HLA-A*11:01 and the frequencies of shared and patient-specific CR ncHLAp. **(B)** Comparison of the expression levels of CR ncHLAp source transcripts across 26 healthy tissues (dGTEx, left heatmap) and the medulloblastoma samples investigated in this study (middle heatmap). The detection of the corresponding CR ncHLAp across the medulloblastoma samples is shown in the right heatmap. The 30 CR ncHLAp with the highest source-transcript expression levels in medulloblastoma were selected for visualization. **(C)** Recurrence of CR ncHLAp across different tumor types. The 30 CR ncHLAp with the highest detection frequencies across tumor samples in the IEAtlas dataset were selected and displayed in the heatmap. HLA-presented peptides detected in tumor samples from the IEAtlas dataset were considered tumor-specific. **(D)** Representative MS/MS spectrum of the CR ncHLAp shown in the top row of the heatmap in (C).

### CR ncHLAp source genes were shared across patients with medulloblastoma

To investigate whether the source genes of CR ncHLAp were shared across patients with medulloblastoma, we identified the source genes of these peptides and assessed their representation across tumor samples. Consistent with previous reports, CR ncHLAp derived from several source genes were detected in two or more medulloblastoma samples (Fig. 4A) [5, 29]. Analysis of 30 selected source genes across the 21 medulloblastoma samples revealed that (i) some source genes gave rise to multiple distinct CR ncHLAp; (ii) most CR ncHLAp originating from shared source genes showed favorable predicted binding to their corresponding HLA allotypes; and (iii) more than half of the analyzed source genes had previously been implicated in tumorigenesis or tumor progression, including TOPBP1 and H1-2 (Fig. 4B, Table 1 and Supplementary Table 1). TOPBP1 has been reported to support the oncogenic activity of mutant p53 and promote tumor progression, highlighting its potential as a therapeutic target in cancer [10]. Linker histone H1-2 has also been shown to contribute to hepatocarcinogenesis by regulating STAT3 activation [44]. The generation and presentation of CR ncHLAp from genes implicated in tumorigenesis may reflect tumor-associated dysregulation of transcription, RNA processing, or translation, although the underlying mechanisms require further investigation. Collectively, these findings demonstrate that CR ncHLAp can arise from source genes recurrently represented across patients with medulloblastoma, including genes implicated in tumor development and progression.

**Fig. 4.**
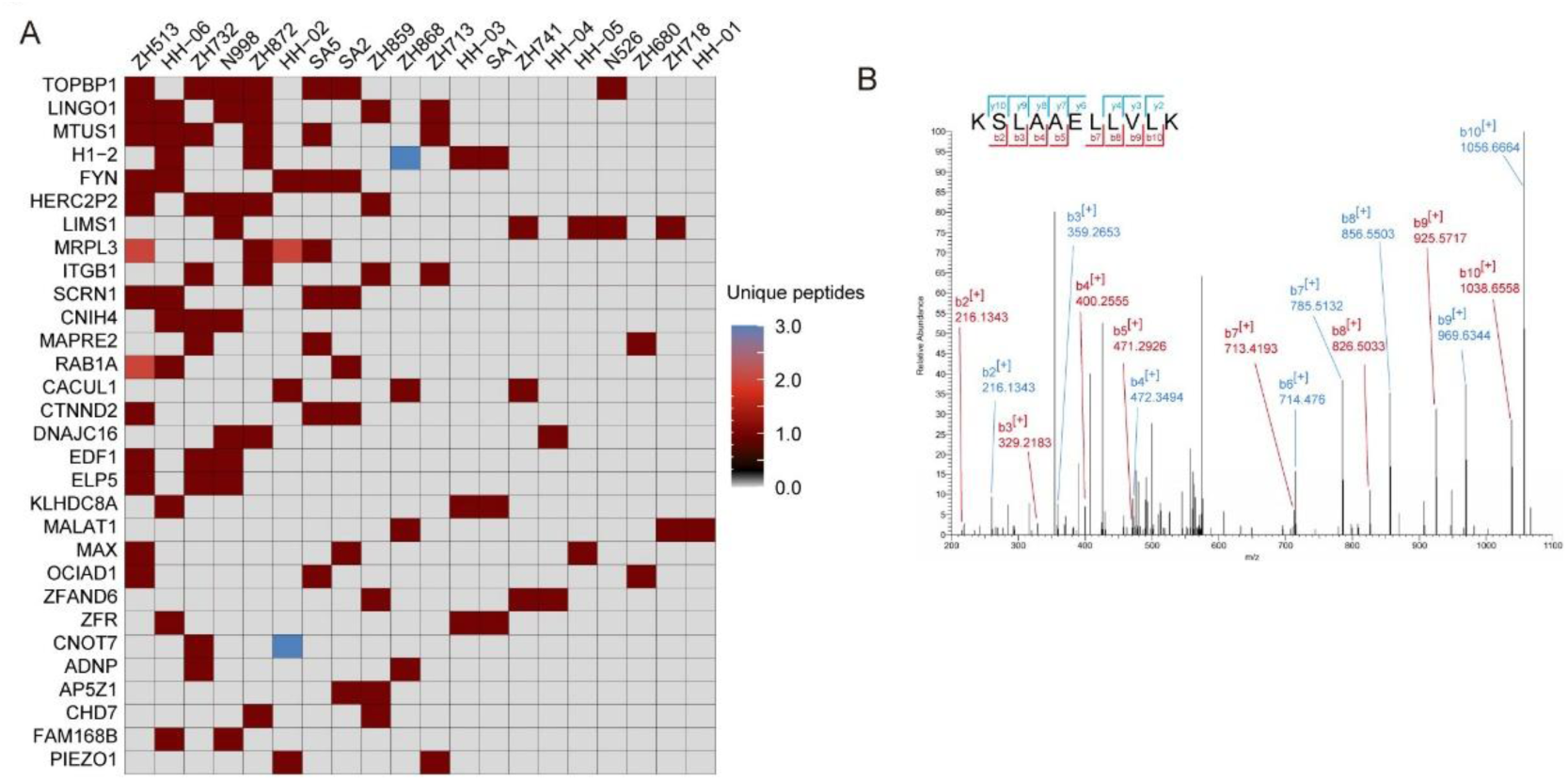
Source genes of CR ncHLAp were shared across patients with medulloblastoma. **(A)** Heatmap showing the distribution of 30 CR ncHLAp source genes across multiple medulloblastoma samples. The number of CR ncHLAp derived from each source gene is also indicated. **(B)** Representative MS/MS spectrum of the CR ncHLAp derived from TOPBP1.

**Table 1.**
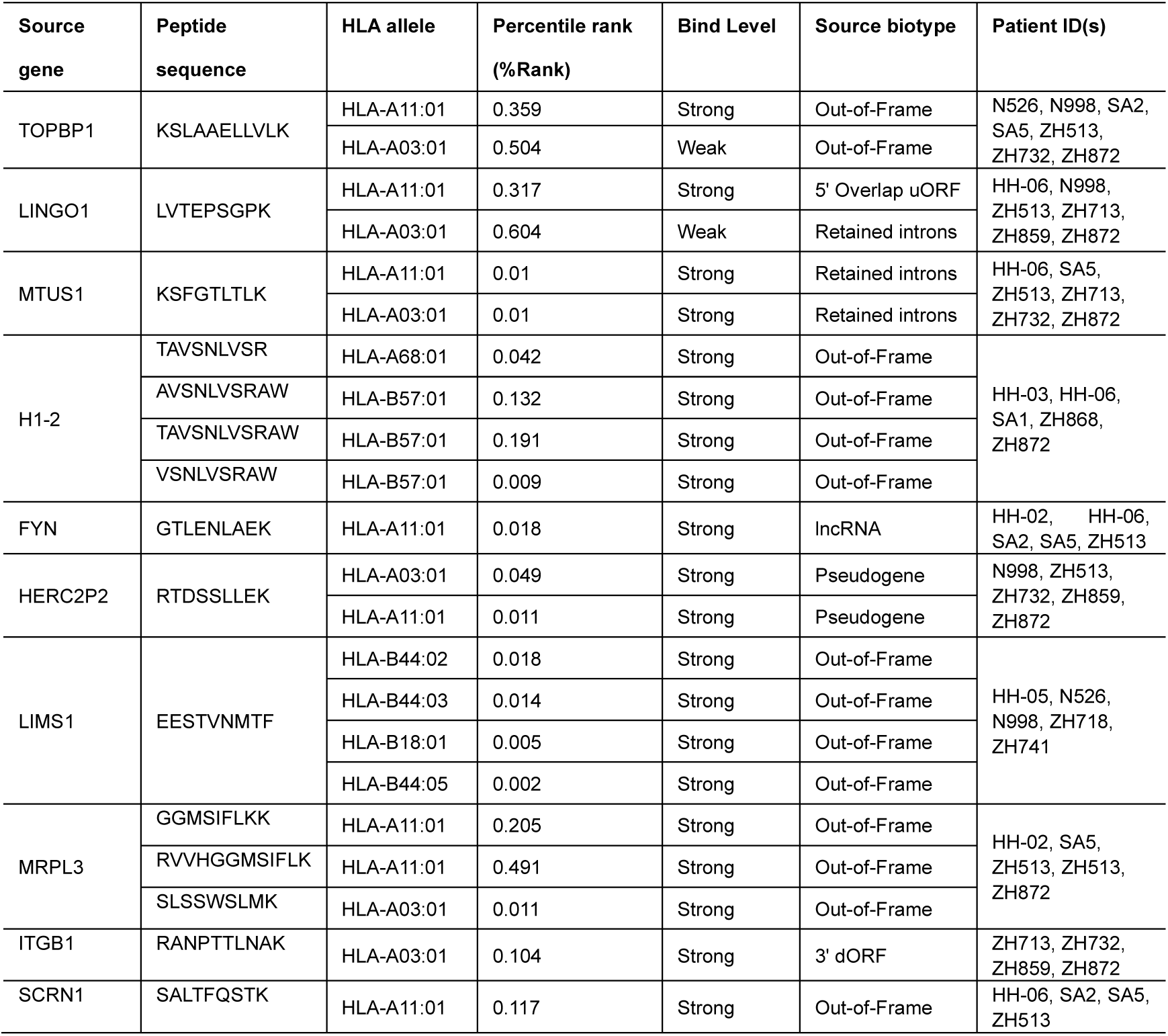
Source genes of representative CR ncHLAp identified in medulloblastoma patients.

### Tumor-specific RNA AS did not correlate with CR ncHLAp burden

In order to explore the potential mechanisms underlying the generation of CR ncHLAp, we performed gene set enrichment analysis (GSEA) to identify biological processes enriched in each medulloblastoma subgroup compared with normal cerebellum samples from GTEx (Supplementary Fig. 3A-3D). As shown in Fig. 5A-5B and Supplementary Fig. 4A-4B, biological processes potentially related to AS, including nucleosome assembly, DNA replication-dependent and -independent chromatin organization, and nuclear mRNA deadenylation dependent decay, were enriched in medulloblastoma subgroups. According to previous reports, nucleosome occupancy and chromatin structure can influence AS [7, 17, 26]. In addition, aberrantly spliced transcripts may be eliminated through RNA surveillance pathways such as nonsense-mediated mRNA decay, which can involve transcript deadenylation [26, 28]. To further investigate the abundance of AS event in medulloblastoma, we analyzed the RNA-seq data using rMATS, with RNA-seq data from GTEx normal cerebellum samples used as the reference. AS events, including alternative 3′ splice site (A3SS), alternative 5′ splice site (A5SS), novel junction, retained intron (RI), and skipped exon (SE), were detected at varying frequencies across patients with medulloblastoma, and nearly half of these events were classified as tumor-specific (Fig. 5C). Among both tumor-enriched and tumor-specific AS events, SE was the most frequent event type and affected the largest numbers of transcripts and corresponding genes. Moreover, multiple transcripts harbored distinct combinations of AS event types (Fig. 5D and Supplementary Fig. 4C). The numbers of tumor-specific RI, A3SS, A5SS, and SE events, as well as the total number of tumor-enriched AS events, were broadly comparable across patients, except for patient 9, who exhibited the fewest tumor-specific SE events and the lowest total number of tumor-enriched AS events (Fig. 5E-5H and Supplementary Fig. 4D). Importantly, no clear association was observed between the number of tumor-specific AS events and the number of CR ncHLAp detected in individual patients. These findings suggest that the overall burden of tumor-specific AS events is unlikely to be a major determinant of CR ncHLAp abundance in medulloblastoma, although they do not exclude the possibility that individual CR ncHLAp may arise from specific AS events.

**Fig. 5.**
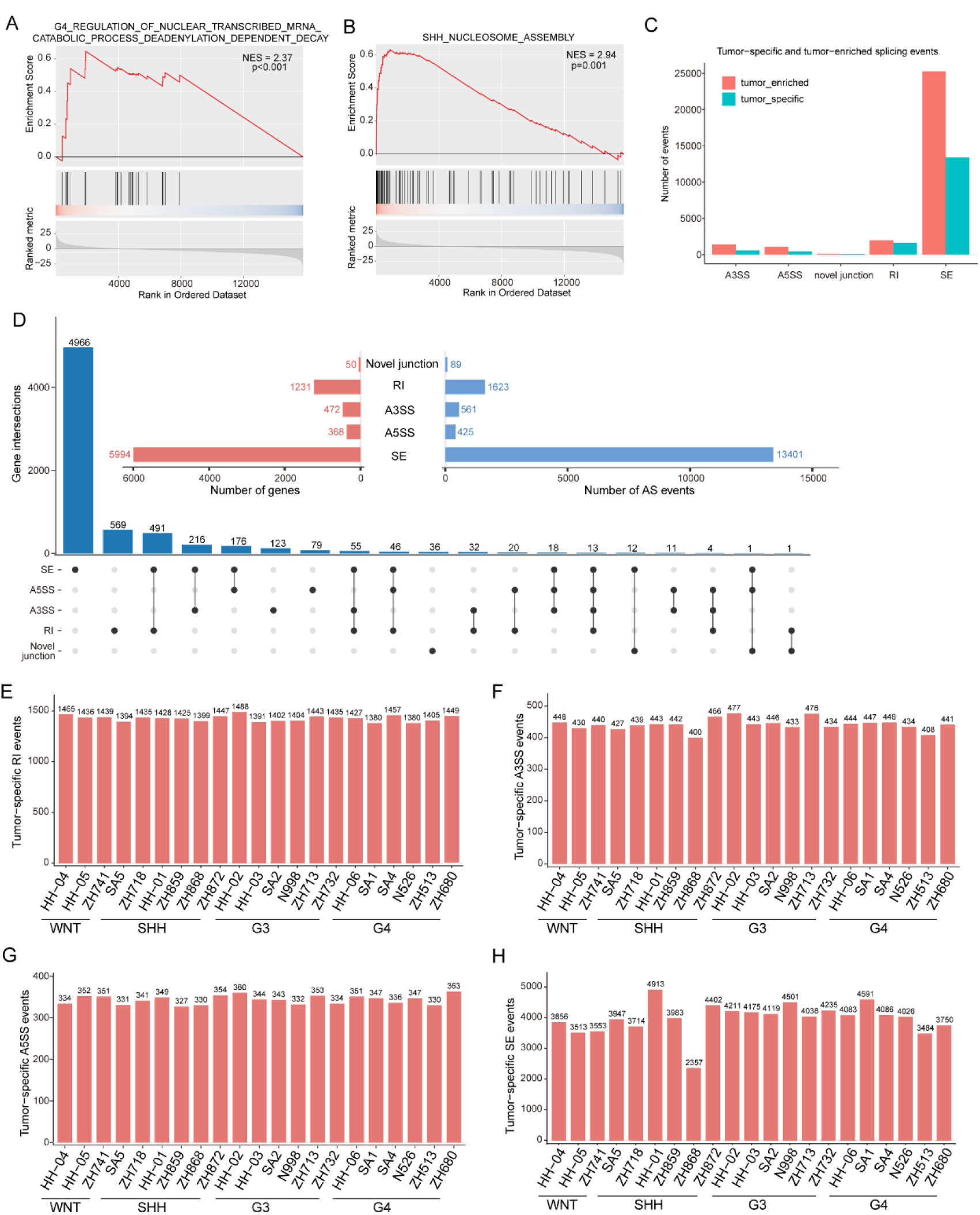
Aberrant AS events in medulloblastoma samples. Representative GSEA plots showing the enrichment of pathways related to deadenylation-dependent mRNA decay **(A)** and nucleosome assembly **(B)** in medulloblastoma relative to normal cerebellum samples from GTEx. NES, normalized enrichment score. **(C)** Numbers of tumor-enriched and tumor-specific transcript variants detected across the medulloblastoma samples, using GTEx normal cerebellum samples as the reference. **(D)** Numbers of genes associated with each category of tumor-specific splicing event, including novel junctions, RI, A3SS, A5SS, and SE. The number of genes intersecting across different AS types is also displayed. RI, intron retention; A3SS, alternative 3′ splice site; SE, skipped exon; A5SS, alternative 5′ splice site. **(E-H)** Numbers of tumor-specific RI (E), A3SS (F), A5SS (G), and SE (H) events detected in each medulloblastoma sample.

## Discussion

By integrating immunopeptidomic and transcriptomic data, we identified 598 CR ncHLAp in 21 medulloblastoma samples, thereby expanding the repertoire of candidate targets for T cell-based immunotherapy in this malignancy. These CR ncHLAp were derived mainly from intron-retaining transcripts, out-of-frame ORFs, and lncRNAs (Fig. 2E). NetMHCpan 4.1 predicted that more than 70% of the CR ncHLAp could bind their corresponding HLA allotypes, whereas DeepImmuno predicted approximately 87% to be immunogenic (Fig. 2H-I), suggesting that these peptides may have potential as therapeutic targets for clinical translation. In addition, some CR ncHLAp were not only shared across multiple medulloblastoma samples but also detected in other cancer types, suggesting that this subset of peptides could be used to design of off-the-shelf multi-epitope cancer vaccines. However, these CR ncHLAp require further functional validation in future studies.

Like some solid tumors, medulloblastoma harbors a low mutation rate, with a median non-synonymous mutation rate of approximately 0.35 mutations per megabase [36, 43]. The availability of mutation-derived neoantigens is limited in tumors with a low mutational burden. Although encouraging clinical responses have been observed in some patients receiving neoantigen vaccines, only 6-11% of predicted neoantigens have been reported to be immunogenic [38, 39, 47]. Therefore, HLA-presented peptides stemmed from mutated sequences alone may provide an insufficient pool of targets for individualized neoantigen-specific immunotherapy in medulloblastoma. Although several TAAs and TSAs, including antigens derived from SNVs, aberrant splice junctions, and gene fusions, have been identified in medulloblastoma, their ability to inhibit tumor growth in appropriate models remains unclear [37, 42]. Taken together, these findings suggest that investigating HLA-presented peptides derived from non-canonical sources may be necessary to broaden the repertoire of targetable antigens for medulloblastoma immunotherapy.

Comprehensive and accurate identification of the HLA ligandome is critical for the development of cancer immunotherapies, including cancer vaccines and adoptive cell therapies. To date, HLA-I immunoprecipitation coupled with mass spectrometry (HLA-I IP-MS) is currently the principal unbiased approach for directly identifying naturally presented HLA-I ligands at the sequence level. However, MS-based identification of ncHLAp typically requires concatenating canonical proteome references with predicted polypeptide sequences derived from non-canonical sources, such as novel unannotated open reading frames and retained introns, thereby inflating the database search space. Under stringent FDR control, searching MS data against an inflated database may decrease the sensitivity of HLA-bound peptide identification, leading to the exclusion of some true ncHLAp [2, 15, 32]. In addition, previous studies have demonstrated that reducing the size of personalized search databases can improve the accuracy of ncHLAp identification and increase the proportion of ncHLAp among the identified peptides [8, 32]. Moreover, canonical HLAp and ncHLAp exhibit distinct score distributions; therefore, pooling them together may increase the error rate in the ncHLAp group. Separate FDR estimation for these two groups is essential to control the rate of false identifications [14]. Therefore, in the present study, we applied a two-stage database search strategy with reference to previously reported workflows [18, 49]. To reduce database complexity and search-space size, raw MS data were first searched against the sample-specific personalized protein databases constructed in this study using a 5% group-specific FDR threshold. Peptide sequences retained after the first-stage search were then used to construct reduced personalized protein databases. In the second-stage search, the raw spectra were searched again against the reduced personalized protein databases using a 1% group-specific FDR threshold.

Using this strategy, we identified 2,298 unique ncHLAp across 21 medulloblastoma samples. Among these peptides, 598 were considered cancer-restricted after stringent filtering using Ribo-seq and immunopeptidomics data from healthy tissues, with an average of 28 CR ncHLAp per patient (Fig. 2A-2B). Most CR ncHLAp were predicted to bind to their corresponding HLA alleles and to be immunogenic (Fig. 2H-2I). Nevertheless, the number of CR ncHLAp identified in the present study was relatively low, which was likely attributable to the limited sensitivity of mass spectrometry and the methodological strategy employed in this study. In addition, although some CR ncHLAp showed no translational evidence in the available healthy-tissue datasets, their source transcripts were detected in healthy tissues (Fig. 3B). As more sensitive analytical approaches become available, it remains possible that a subset of CR ncHLAp may show translational evidence in normal tissues. However, the finding suggests that our filtering pipeline was sufficiently sensitive to identify and exclude 99.04% of the canonical HLAp whose source ORFs showed evidence of translation in normal tissues (Fig. 2C). In addition, paired immunopeptidomic and transcriptomic datasets from developing normal cerebellum would represent the ideal control for identifying CR ncHLAp in medulloblastoma. However, obtaining normal cerebellar tissue from developing individuals through surgical resection is ethically infeasible. Because normal cerebellar transcriptomic data are unavailable in dGTEx, we used transcriptomic data from normal cerebellum in GTEx as the control to identify aberrant AS events in medulloblastoma. Notably, most of the CR ncHLAp identified in this study were recurrently detected across tumors in the IEAtlas dataset [4], supporting the robustness of our analysis.

Our study demonstrated that some CR ncHLAp were not only shared among multiple medulloblastoma samples but also recurrently detected in other cancer types, suggesting that common oncogenic processes may contribute to their generation (Fig. 3A-3C). For example, oncogenic RAS signaling has been reported to promote translation initiation and increase translational flux [12]; c-MYC can change translation initiation sites and mRNA translation efficiency [40]; and SF3B1 mutations, which frequently occur in cancer, can induce aberrant 3′ splice-site selection and promote tumor-specific splicing alterations [11]. In the present study, several tumorigenesis-associated genes, such as TOPBP1 and H1-2, were found to give rise to CR ncHLAp (Fig. 4A), suggesting that cancer-associated genes may serve as important sources of CR ncHLAp. In addition, GSEA revealed the enrichment of biological processes related to chromatin organization, nucleosome assembly, and deadenylation-dependent mRNA decay in medulloblastoma (Fig. 5A-5B and Supplementary Fig. 4A-4B). Given that alterations in chromatin structure and nucleosome occupancy can reshape splicing patterns, that many alternatively spliced transcripts containing premature termination codons (PTCs) undergo nonsense-mediated mRNA decay, and that deadenylation represents an initial step in most mRNA decay pathways [6, 16, 31], we hypothesized that aberrant AS may contribute to the generation of CR ncHLAp in medulloblastoma. Therefore, we analyzed cancer-specific AS events, including RI, A3SS, A5SS, and SE, in individual patients (Fig. 5C-5H). Although a substantial cancer-specific AS events were identified, and multiple types of AS events were detected in many genes or transcripts, no significant association was observed between the number of cancer-specific AS events and the number of CR ncHLAp identified per patient. These findings may be partly attributable to limitations of our analytical approach. Further investigation is therefore required to clarify the mechanisms underlying the generation of CR ncHLAp in medulloblastoma.

### Limitations of the study

Although this study mapped the CR ncHLAp landscape of medulloblastoma and highlighted the potential of these peptides as candidate targets for T cell-based immunotherapy, several limitations should be acknowledged. First, *in vitro* and *in vivo* experiments assessing the immunogenicity of these CR ncHLAp were not performed and should be conducted in future studies. Second, paired immunopeptidomic and transcriptomic data from cerebellar tissues adjacent to tumors or from healthy developing cerebellar tissues, which could serve as negative controls, were unavailable in this study. Third, the nomination of CR ncHLAp relied on limited immunopeptidomic and Ribo-seq datasets from normal tissues. Finally, the mechanisms underlying the generation of CR ncHLAp in medulloblastoma remain unclear and warrant further investigation.

### Additional files

#### Additional file 1 (.docx)

Supplementary Figures. This file contains Supplementary Figs. 1-4 and their corresponding legends.

#### Additional file 2 (.docx)

Supplementary Table. This file contains Supplementary Table 1, which lists the source genes of representative CR ncHLAp identified in medulloblastoma samples.

## Declarations

### Ethics approval and consent to participate

Not applicable.

### Consent for publication

Not applicable.

### Data availability

The publicly available RNA-seq data analyzed in this study were obtained from the Gene Expression Omnibus (GEO; accession number GSE238195; https://www.ncbi.nlm.nih.gov/geo/query/acc.cgi?acc=GSE238195). The mass spectrometry data used in this work were obtained from the PRIDE Archive and are publicly available under accession number PXD043770 (https://www.ebi.ac.uk/pride/archive/projects/PXD043770). The data supporting the findings of this study are included in the article and its supplementary figures. Additional derived data are available from the corresponding author upon request.

## Authors’ contributions

Study design: A. H. Acquisition and analysis of the data: N. Z. and Y. X. Creation of new code: W. M. Drafting of the initial manuscript: N. Z. Revision of the manuscript: N. Z., Y. X., and W. M. All authors have read and approved the contents of final manuscript.

## Supporting information

Supplementary table 1. Source genes of representative CR ncHLAp identified in medulloblastoma samples

## Acknowledgments

This work was supported by EnCureGen Pharma Guangzhou Limited.

