## Supplementary Figure 1-4 for "Integrated immunopeptidomics and transcriptomics reveal a cancer-restricted non-canonical HLA class I ligandome in medulloblastoma"

Supplementary Fig. 1

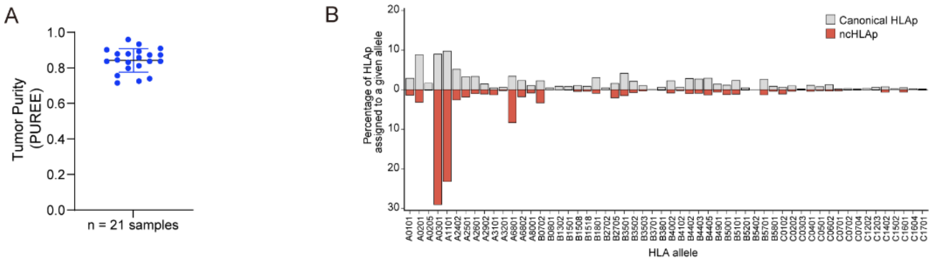


**Supplementary Fig. 1 Estimated tumor purity and predicted HLA binding of peptides identified in medulloblastoma samples. (A)** The tumor purity of the analyzed medulloblastoma samples was estimated using the PUREE algorithm. **(B)** Percentage of canonical HLAp or ncHLAp predicted to bind a given allele. Only peptides predicted to bind at least one patient-matched HLA allele were included in this analysis.

Supplementary Fig. 2

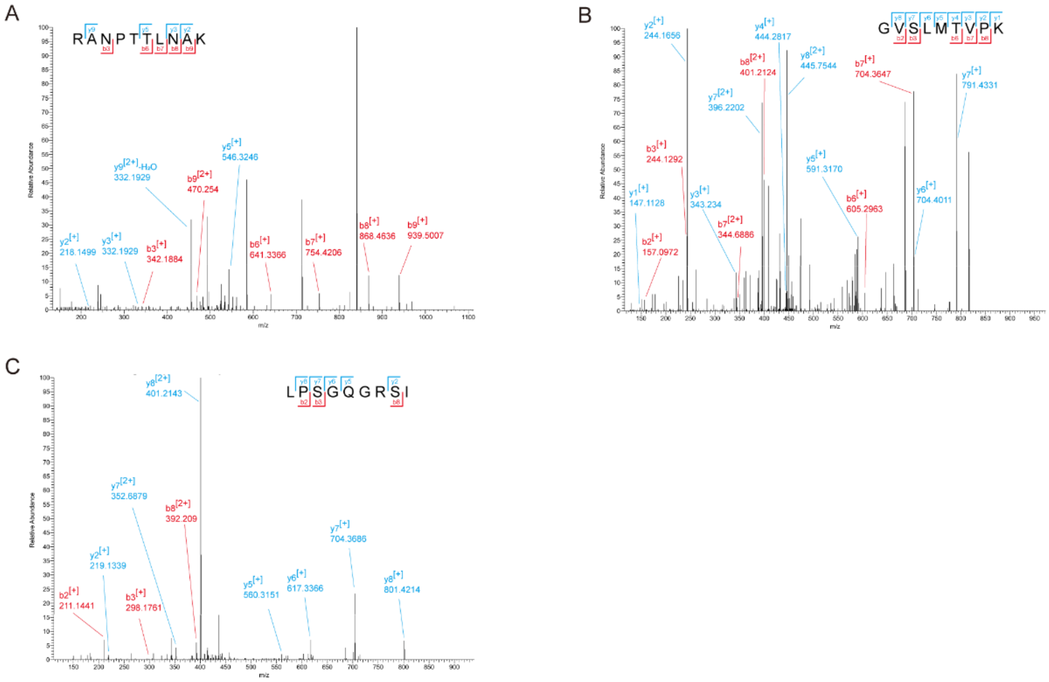


**Supplementary Fig. 2 Representative MS/MS spectra of CR ncHLAp. (A-C)** MS/MS spectra of the three CR ncHLAp with the most favorable predicted HLA-binding ranks for their corresponding HLA allotypes.

Supplementary Fig. 3

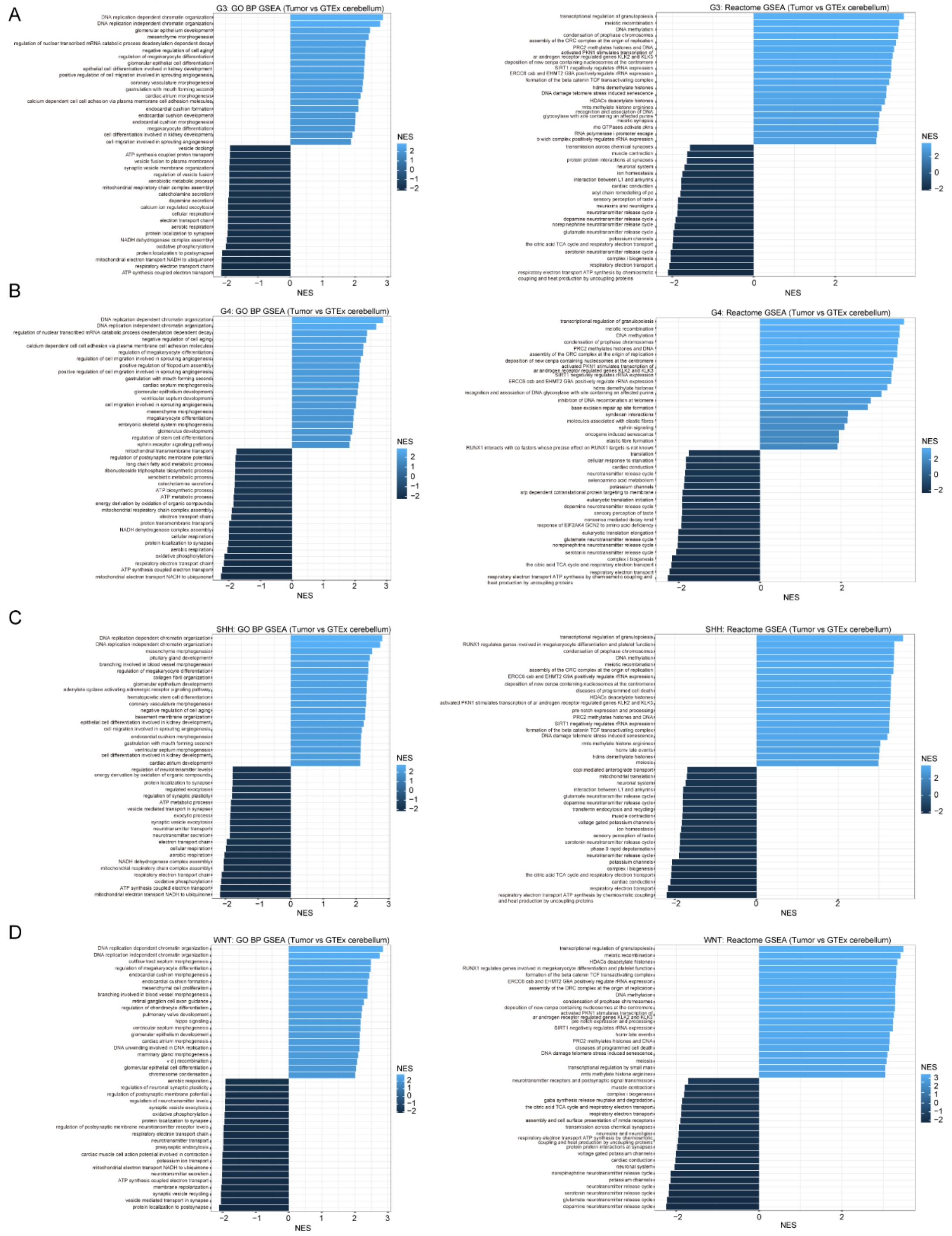


**Supplementary Fig. 3 GSEA for medulloblastoma subgroups. (A-D)** Bar plots depicting Gene Ontology (GO) Biological Process (BP) terms and Reactome pathways exhibiting significant positive or negative enrichment in G3 (A), G4 (B), SHH (C), and WNT (D) medulloblastoma samples relative to normal cerebellum samples from GTEx.

Supplementary Fig. 4

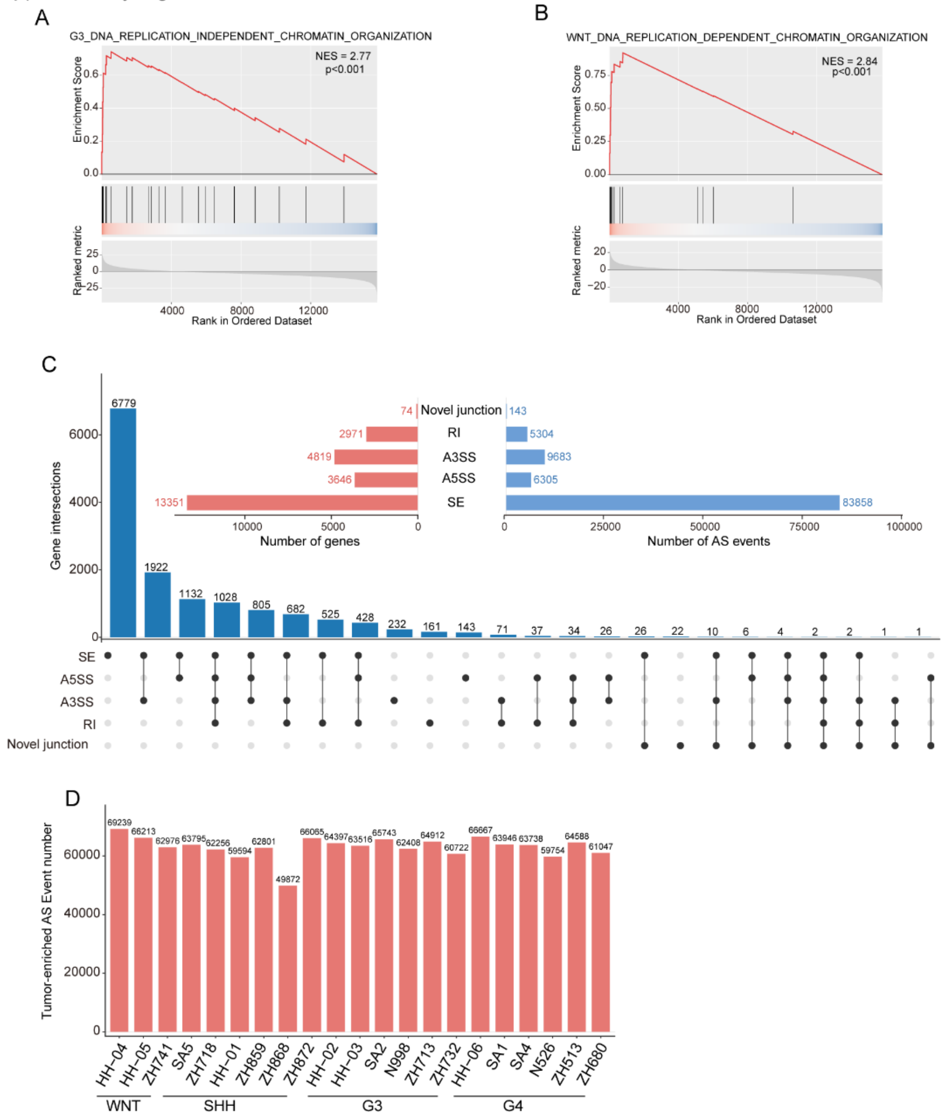


**Supplementary Fig. 4 Abnormal AS events in medulloblastoma samples.** Representative GSEA plots showing the enrichment of DNA replication-independent chromatin organization **(A)** and DNA replication-dependent chromatin organization **(B)** in medulloblastoma samples relative to normal cerebellum samples from GTEx. **(C)** Numbers of tumor-enriched AS events and associated genes for each AS event type. The numbers of genes shared among different combinations of AS event types are also shown. **(D)** Total numbers of tumor-enriched AS events detected in individual medulloblastoma samples.
