## Supplementary table 1. Source genes of representative CR ncHLAp identified in medulloblastoma samples for "Integrated immunopeptidomics and transcriptomics reveal a cancer-restricted non-canonical HLA class I ligandome in medulloblastoma"

Supplementary table 1. Source genes of representative CR ncHLAp identified in medulloblastoma patients.

| Source gene | Peptide sequence | HLA allele | Percentile rank (%Rank) | Bind Level | Source biotype | Patient ID(s) |
|---|---|---|---|---|---|---|
| CNIH4 | ISYDLSFDK | HLA-A11:01 | 0.144 | Strong | Out-of-Frame | HH-06, N998, ZH732 |
| | KRNGEMVGK | HLA-A03:01 | 3.410 | | 3' dORF | |
| MAPRE2 | WRSMCILPR | HLA-B27:05 | 1.236 | Weak | 5' Overlap uORF | SA5, ZH680, ZH732 |
| | GSVPTVGEW | HLA-B58:01 | 0.019 | Strong | 5' Overlap uORF | |
| RAB1A | ATSAQLVWISK | HLA-A11:01 | 0.087 | Strong | Out-of-Frame | HH-06, SA2, ZH513, ZH513 |
| | SAQLVWISK | HLA-A11:01 | 0.207 | Strong | Out-of-Frame | |
| CACUL1 | TEMSAILLL | HLA-B50:01 | 0.297 | Strong | Retained introns | HH-02, ZH741, ZH868 |
| | | HLA-B44:05 | 0.009 | Strong | Retained introns | |
| | | HLA-B44:02 | 0.129 | Strong | Retained introns | |
| CTNND2 | SVAVTRWSK | HLA-A11:01 | 0.063 | Strong | Out-of-Frame | SA2, SA5, ZH513 |
| DNAJC16 | SIASAISMK | HLA-A03:01 | 0.030 | Strong | 5' Overlap uORF | HH-04, N998, ZH872 |
| EDF1 | RTNNILLPR | HLA-A03:01 | 0.107 | Strong | Out-of-Frame | N998, ZH513, ZH732 |
| | | HLA-A11:01 | 0.031 | Strong | Out-of-Frame | |
| ELP5 | ALWELSAALLR | HLA-A03:01 | 1.089 | Weak | 3' dORF | N998, ZH513, ZH732 |
| KLHDC8A | EVAIGAVGAGAK | HLA-A68:01 | 1.032 | Weak | 5' uORF | HH-03, HH-06, SA1 |
| MALAT1 | IMGEFRTEV | HLA-A02:01 | 0.174 | Strong | lncRNA | HH-01, ZH718, ZH868 |
| MAX | SVLQSDRFIK | HLA-A11:01 | 0.255 | Strong | 3' dORF | HH-05, SA2, ZH513 |
| OCIAD1 | MRVGLLLSL | HLA-B27:05 | 0.020 | Strong | Retained introns | SA5, ZH513, ZH680 |
| ZFAND6 | LFSAARFLF | HLA-A24:02 | 0.132 | Strong | 3' dORF | HH-04, ZH741, ZH859 |
| ZFR | TTLDSPTAGR | HLA-A68:01 | 0.037 | Strong | Retained introns | HH-03, HH-06, SA1 |
| CNOT7 | ASSAIQVHK | HLA-A11:01 | 0.008 | Strong | 5' uORF | HH-02, ZH732 |
| | SASSAIQVHK | HLA-A11:01 | 0.159 | Strong | 5' uORF | |
| | SSAIQVHK | HLA-A11:01 | 1.173 | Weak | 5' uORF | |
| | RSASSAIQVHK | HLA-A03:01 | 0.174 | Strong | 5' uORF | |
| ADNP | RLWKHTLKY | HLA-A80:01 | 0.001 | Strong | Out-of-Frame | ZH732, ZH868 |
| | GLSDKTFLL | HLA-A02:01 | 0.027 | Strong | Out-of-Frame | |
| AP5Z1 | PLTLCQRCVS | HLA-C07:01 | 59.592 | | Retained introns | SA2, ZH859 |
| | HPARDVPL | HLA-B35:02 | 0.250 | Strong | Retained introns | |
| CHD7 | IYSGTFLYL | HLA-A24:02 | 0.019 | Strong | Retained introns | ZH859, ZH872 |

| | | | | | | |
|---|---|---|---|---|---|---|
| | GSLNLSMLK | HLA-A03:01 | 0.139 | Strong | Retained introns | |
| FAM168B | VRLLAGFIV | HLA-B52:01 | 11.334 | | 3' dORF | HH-06, N998 |
| | PPAGLCHRTPPP | HLA-C16:01 | 34.158 | | Out-of-Frame | |
| PIEZO1 | CCPTVLALT | HLA-A02:05 | 32.240 | | Retained introns | HH-02, ZH713 |
| | ASVGGARPARAWL | HLA-C12:03 | 23.815 | | Retained introns | |
